# Re-evaluating the relationship between female social bonds and infant survival in wild baboons

**DOI:** 10.1101/2024.08.20.608854

**Authors:** Maria J.A. Creighton, Brian A. Lerch, Elizabeth C. Lange, Joan B. Silk, Jenny Tung, Elizabeth A. Archie, Susan C. Alberts

## Abstract

Over the past few decades studies have provided strong evidence that the robust links between the social environment, health, and survival found in humans also extend to non-human social animals. A number of these studies emphasize the early life origins of these effects. For example, in several social mammals, more socially engaged mothers have infants with higher rates of survival compared to less socially engaged mothers, suggesting that positive maternal social relationships causally improve offspring survival. Here we show that the relationship between infant survival and maternal sociality is confounded by previously underappreciated variation in female social behavior linked to changes in reproductive state and the presence of a live infant. Using data from a population of wild baboons living in the Amboseli basin of Kenya – a population where high levels of maternal sociality have previously been linked to improved infant survival – we find that infant- and reproductive state-dependent changes in female social behavior drive a statistically significant relationship between maternal sociality and infant survival. After accounting for these state-dependent changes in social behavior, maternal sociality is no longer positively associated with infant survival in this population. Our results emphasize the importance of considering multiple explanatory pathways—including third-variable effects—when studying the social determinants of health in natural populations.

## INTRODUCTION

In social species, social interactions play a major role in determining an animal’s access to resources (Wenzel & Pickering, 1991; Ranta et al., 1993; Wright et al., 2001; Silk, 2007), exposure to pathogens (Kappeler et al., 2015), protection against predators (Sterck et al., 1997; Wright et al., 2001; Krause & Ruxton, 2002; Silk, 2007), and adoption of novel behaviors (Burkart, 2017). Consistent with this idea, the social environment has been linked to health and survival outcomes in a wide variety of social mammals, including humans (Silk et al., 2010; Archie et al., 2014; Brent et al., 2017; Ellis et al., 2019; Campos et al., 2020; Snyder-Mackler et al., 2020). For the dependent young of many species, the social environment is primarily determined by the social bonds of their mothers. For this reason, maternal social traits have been proposed to influence offspring survival, with evidence supporting this link presented in non-human primates (Silk et al., 2003a; Silk et al., 2009; Kalbitzer et al., 2017; McFarland et al., 2017; Schneider-Crease et al., 2022; Blersch et al., 2023), dolphins (Frère et al., 2010), sheep (Vander Wal et al., 2015), and horses (Cameron et al., 2009).

Because of their long period of maternal dependence and close evolutionary relationship to humans, non-human primates have been a particular focus of studies linking maternal sociality to offspring survival. In the Amboseli baboon population of southern Kenya (an admixed population of yellow, *Papio cynocephalus*, and anubis, *P. anubis*, baboons) females that have the strongest social bonds show the highest relative infant survival over their lifetime (Silk et al., 2003a). In a long-term study of chacma baboons in Botswana (*P. ursinus*), offspring of females with stronger social bonds also live significantly longer lives (Silk et al., 2009). Similarly, in the De Hoop chacma baboons of South Africa, baboon infants whose mothers have many weak social bonds are also more likely to survive the first 12-months of life than infants whose mothers have fewer weak social bonds (McFarland et al., 2017) and evidence in vervet monkeys (*Chlorocebus pygerythrus*) suggests infant survival increases with the number of maternal spatial partners (Blersch et al., 2023). These studies suggest a positive role of maternal sociality in bolstering offspring survival. However, Schneider-Crease et al. (2022) recently revealed no significant relationship between maternal sociality and offspring survival in kinda baboons (*P. kindae*) and in white-faced capuchins (*Cebus imitator*), the offspring of highly social females exhibit higher survivorship than those of less social females during socially stable periods, but lower survivorship during the less stable periods surrounding alpha male turnover (Kalbitzer et al., 2017). Therefore, associations between maternal sociality and offspring survival within primates appear to vary as a function of species, population, and/or prevailing demographic conditions.

Notably, all analyses of the relationship between maternal social behavior and infant survival face a key challenge: the amount of time a mother has a living infant may itself drive patterns of female sociality. For example, in a number of primate species, conspecific adult females are attracted to young infants (Seyfarth, 1976; Altmann, 1980; Small, 1982; Silk et al., 2003b; Tiddi et al., 2010; Dunayer & Berman, 2018). This effect could cause mothers with surviving infants to appear more social than mothers whose infants die, simply because they have a socially attractive infant for a longer time period (Barrett et al., 2007). Under this scenario, infant survival drives estimates of female sociality, instead of female sociality driving infant survival, providing an explanation that is plausible regardless of whether maternal sociality is measured over the lifetime (e.g., Silk et al., 2003a) or over fixed yearly intervals (e.g., Kalbitzer et al., 2017, Silk et al., 2009; McFarland et al., 2017; Schneider-Crease et al., 2022). Similarly, the relationships females have with males may depend on patterns in infant survival. In many primates the death of an infant is followed by the rapid resumption of sexual cycling (the state during which grooming and proximity to males peak) and males may also be socially attracted to young neonates (e.g., Nguyen et al., 2009; Baniel et al., 2016). Some approaches to avoiding infant-dependent variation in female social relationships in such analyses include discarding maternal social interactions that occur when infants are very young (e.g., less than 100 days: Silk et al., 2009) or measuring maternal sociality before an infant is born (Blersch et al., 2023). However, these approach have limitations. The first approach does not elimate infant-dependent variation in maternal sociality that occurs after the discarded time window (e.g., cycling resumption following an infant’s death). Moreover, both approaches by design eliminate the social interactions that are likely to be the most consequential for infant survival, possibly obscuring the true causal effects of maternal social relationships on offspring survival.

Here, we directly address the potential influence of infant-driven variation in maternal social behavior by measuring and adjusting for the magnitude of this potential confound in a study of the baboons living in the Amboseli basin of southern Kenya – a population where higher levels of maternal lifetime sociality have previously been linked to improved infant survival (Silk et al., 2003a). We use a previously established measure of maternal social behavior (maternal social connectedness, or SCI; Archie et al., 2014, see Methods) to demonstrate that a female’s social interactions with both adult females and adult males are strongly dependent upon her reproductive state along with the age and survival status of any current infant. Next, we demonstrate how the relationships between a female’s reproductive state, infant age/presence, and her rates of social interactions produce statistically significant associations between maternal SCI and infant survival that can change in both direction and magnitude depending upon the time interval over which SCI is measured. After accounting for infant- and reproductive state-driven variation in maternal sociality, we find no compelling evidence that sociality positively predicts infant survival in the present analysis. Finally, we demonstrate that the same confounds attenuate the original effect reported in Silk et al. (2003a) and present some post-hoc analyses assessing the eco-evolutionary consequences of a trade-off between maternal sociality’s effect on adult versus offspring survival.

## RESULTS

### MATERNAL SOCIAL BEHAVIOR STRONGLY DEPENDS ON REPRODUCTIVE STATE AND INFANT PRESENCE

Social connectedness of adult female baboons to other females (SCI-F, a non-dyadic measure of the quantity/overall rate of grooming, the primary affiliative social behavior in baboons) tends to peak for mothers with infants in the first one to six months of life (Fig. 1A; Table S1). This pattern is consistent with the idea that having a young living infant increases a mother’s social interactions with females e.g., by attracting other females to groom with her. Females do not experience this peak in grooming interactions if their infants die in the first year of life (Fig. S1A).

**Figure 1:**
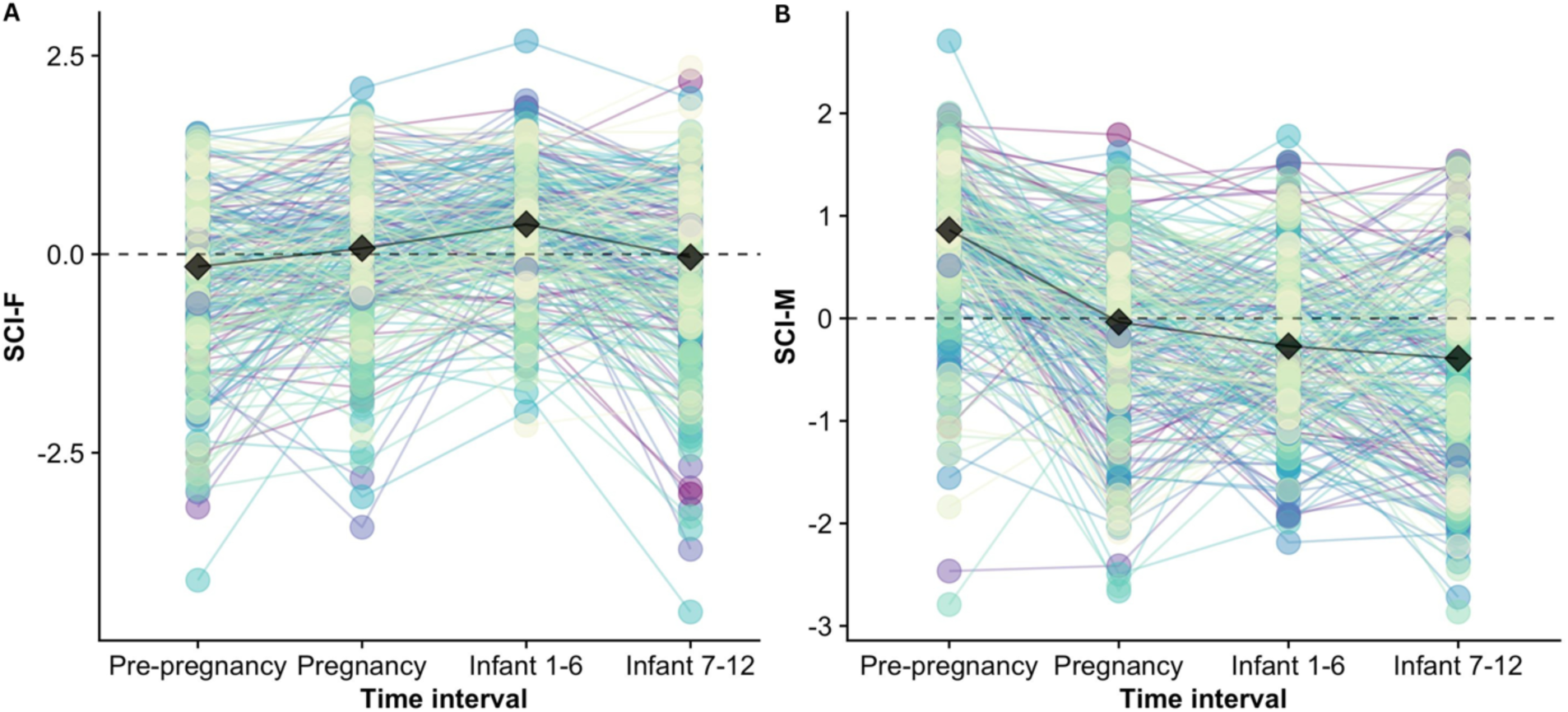
Comparison of A) social connectedness to females (SCI-F) and B) social connectedness to males (SCI-M) for mothers with infants who survived to one year across four six-month time intervals (six months prior to pregnancy, pregnancy, one to six months following birth, and seven to 12 months following birth). Black diamonds show the median for each time interval. Each series of connected dots represents a single infant ID for 257 unique infant births (n=146 unique mothers) where behavioral data for the mother was available for all four time intervals and the mother did not have a young infant (less than six months old) from a previous pregnancy during the pre-pregnancy period.

Social connectedness to males (SCI-M) peaks during the pre-pregnancy period when females are cycling, and steadily declines from pregnancy through the first year of a surviving infant’s life (Fig. 1B; Table S2). If an infant dies, the mother’s connectedness to males returns to pre-pregnancy levels as she resumes cycling (Fig. S1B).

Associations between the presence of a live infant, reproductive state, and maternal SCI could confound an apparent effect of maternal sociality on infant survival. The same concern affects analyses using alternative measures of social integration such as the dyadic sociality index (DSI; Silk et al., 2006; Silk et al., 2013) and the composite sociality index (CSI, which combines data on grooming and proximity to individuals of both sexes; Silk et al., 2003a), which are also affected by reproductive state and infant presence (albeit less so for CSI; Figs. S2 and S3). These patterns indicate that the association between infants, reproductive state, and maternal sociality are not unique to SCI, our specific social measure of interest.

### THE RELATIONSHIP BETWEEN MATERNAL SOCIALITY AND INFANT SURVIVAL VARIES DEPENDING ON WHEN MATERNAL SOCIALITY IS MEASURED

After confirming that mothers experience patterns of social behavior that depend on infants and reproductive state, we investigated the effects of this relationship on the association between maternal social behavior and infant survival. To do so, we analyzed the link between the probability of infant mortality within the first year of life and, in separate models, social connectedness to females (SCI-F) and social connectedness to males (SCI-M) measured during four different time intervals. These time intervals included (i) the fixed six-month window after an infant’s conception (i.e., the pregnancy period in baboons, where gestation lasts an average of six months; Altmann et al., 1977), (ii) the fixed six-month window beginning with the infant’s live birth, (iii) the fixed six-month window beginning seven months after an infant’s birth, and (iv) a ‘shifting time window’ that represents the six months prior to each infant’s death or – if it survived – its first birthday. The shifting time window was designed to exclude any maternal social behavior occurring after (and possibly as a result of) an infant’s death while also capturing key parts of the infant’s life which were likely critical to its survival. The other time intervals occur over the same fixed time period relative to birth for all mothers, regardless of infant outcome, and thus more closely parallel approaches taken in previous studies connecting maternal sociality to infant survival (e.g. Silk et al., 2009; McFarland et al., 2017; Blersch et al., 2023).

We used binomial Generalized Linear Models (GLMs) implemented in R to test the relationship between SCI-F and SCI-M measured over each of the four time intervals and infant survival, controlling for other variables that could influence infant survival (maternal social rank, age, parity, group size, and sexual receptivity; see Methods). In these models the outcome was scored as 1 (if the infant died within one year) or 0 (if the infant survived). The total number of infants included in these analyses varied between 824 and 923 depending on the time interval of interest. This variation in sample sizes was a result of gaps in behavioral data for some mothers during some time intervals when behavioral sampling was constrained (e.g., if groups ranged outside of the study area or demographic events decreased sampling opportunities; see sample sizes reported in Tables S3 to S10). In all data sets, approximately 20% of infants died before their first birthday (range 19 – 24%). For example, in the shifting time window case, 873 infants were included in the analysis, 203 of whom died before reaching age one.

The relationship between SCI-F and infant survival varied dramatically in direction and magnitude depending when SCI-F was measured. SCI-F showed no statistically significant relationship with infant survival when measured over the pregnancy period (coefficient=−0.125; odds ratio (OR)=0.882; p=0.109; Fig. 2; Table S3) or seven to 12 months after an infant’s birth (coef=0.109; OR=1.116; p=0.240; Fig. 2; Table S5). However, when SCI-F was measured over the fixed six-month window directly following a live birth, SCI-F positively predicted survival: infants born to mothers with higher SCI-F experienced lower mortality rates compared to infants born to mothers with a lower SCI-F (coef=−0.208; OR=0.812; p=0.013; Fig. 2; Table S4). In contrast, when measured over the shifting time window, which excludes all time periods after an infant’s death, higher SCI-F negatively predicted survival: infants born to mothers with higher SCI-F experienced higher mortality rates compared to infants born to mothers with a lower SCI-F (coef=0.374; OR=1.454; p=<0.001; Fig. 2; Table S6).

**Figure 2:**
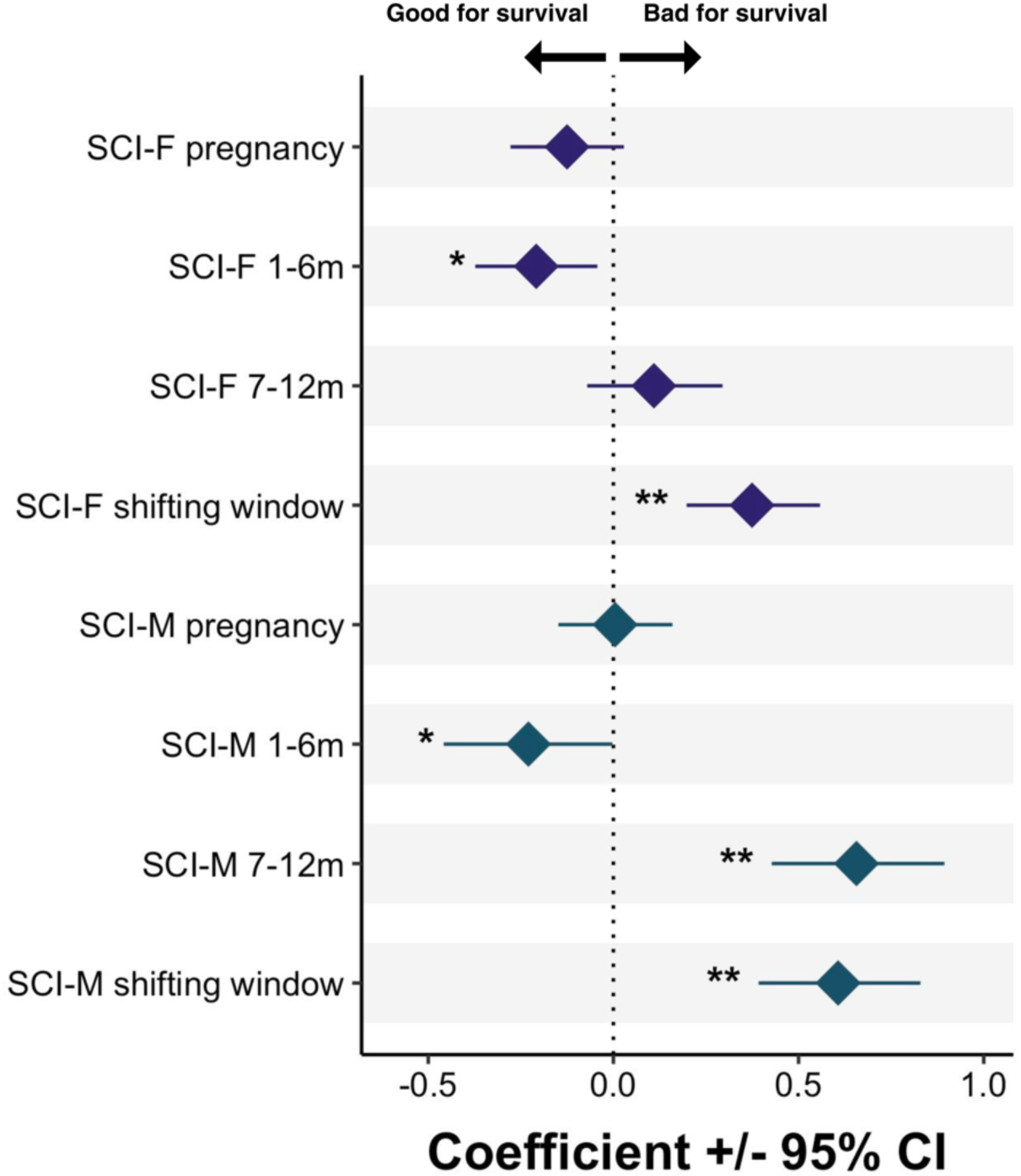
Infant survival as a function of maternal SCI-F and SCI-M measured over different time intervals relative to infant birth; a coefficient (i.e., the natural logarithm of the odds ratio) = 0 (shown by vertical dashed line) indicates no effect of the social measure of interest on infant survival. See Tables S3 to S10 for full model results and sample sizes. +p≤0.1; * p<0.05; ** p<0.01

These contrasting results are consistent with what should be expected if adult females’ attraction to infants, and hence their mothers (Fig. 1A) explain observed associations between infant survival and maternal sociality. Specifically, measuring SCI-F during the fixed six-month time window after birth means that females whose infants survive the entire six-month period appear more social than females whose infants die during that period. Using a shifting time window, however, would cause females whose infants die to appear more social than females whose infants survive because in cases of infant death the mother’s SCI-F tends to be measured earlier in the infant’s life, when infants are most likely to attract social attention.

Similar to SCI-F, the link between SCI-M and infant mortality depended upon the time interval over which SCI-M was measured (Fig. 2). Specifically, SCI-M did not predict infant survival when measured during pregnancy (coef=0.005; OR=1.005; p=0.952; Fig. 2; Table S7). When measured during the six months following birth, higher SCI-M was associated with somewhat lower infant mortality (coef=−0.229; OR=0.795; p=0.048; Fig. 2; Table S8). On the other hand, when measured during the seven to 12 months following birth or using the shifting time window, higher SCI-M was associated with considerably higher infant mortality (coef=0.656; OR=1.928; p=<0.001; Fig. 2; Table S9 and coef=0.607; OR=1.834; p=<0.001; Fig. 2; Table S10, respectively).

As with the SCI-F results, results with SCI-M were consistent with the idea that the reproductive state-dependent nature of female relationships with males affects the apparent relationship between SCI-M and infant survival. Specifically, high levels of sociality with males may be negatively associated with infant survival when measured seven to 12 months after birth because mothers with surviving infants of this age typically have not yet resumed sexual cycling (Gesquiere et al., 2018) and are less social with adult males than females in any other reproductive state (Fig. 1B). In contrast, mothers whose infants die resume sexual cycling and return to their pre-pregnancy levels of social interactions with adult males soon after infant death, resulting in high levels of social interactions with males seven to 12 months after their (non-surviving) infants’ births (Fig. S1B). Similarly, when using the shifting time window, SCI-M for mothers whose infants survive is measured when those mothers are least likely to be interacting with adult males (i.e., when their infants are older but the mothers are not yet cycling).

### INFANT-DEPENDENT AND REPRODUCTIVE STATE-DEPENDENT TRENDS IN MATERNAL SOCIAL BEHAVIOR EXPLAIN THE RELATIONSHIP BETWEEN INFANT SURVIVAL AND MATERNAL SOCIALITY MEASURED OVER MULTIPLE TIME INTERVALS

To determine whether infant-dependent and reproductive state-dependent changes in female social behavior could completely account for the associations between maternal sociality and infant survival in our observed data sets, we used a data randomization procedure to force independence between maternal sociality and infant survival. In this data randomization (hereafter the ‘Randomization of SCI Values’), maternal SCI-F and SCI-M values were assigned randomly to the infant outcomes used in our actual analyses, removing any possible influence of maternal sociality on infant survival. These randomly assigned SCI values were then changed systematically over time, as a function of time since infant’s birth, depending only on whether and when an infant died (see further explanation below). If the confounds we described above were sufficient to account for the observed links between maternal social behavior and infant survival, then the randomized SCI data should drive effects comparable to those obtained with the observed data.

To randomize SCI-F values, we randomly sampled SCI-F trajectories, with replacement, from the set of trajectories shown in Figure 1A (where infant death never occurred), and matched them one-by-one to infant-mother pairs in the true data set (randomization procedure visualized in Fig. S4). For all infants in the true data set we directly substituted the randomly matched SCI-F value during pregnancy for the real SCI-F value during pregnancy. For cases in the real data set where the infant survived to 12 months of age, we also directly substituted the randomly matched SCI-F value in the first six-month interval and the seven to twelve month interval following birth. For cases in the real data where the infant died, we followed the same procedure as for surviving infants with one exception: for all months following the infant’s death we assigned the maternal SCI-F value for pre-pregnancy from the randomly sampled SCI-F trajectory (recall that a female’s pre-pregnancy SCI-F values closely match her SCI-F values after the loss of her infant, Fig. S1). We repeated this procedure across all observed infant outcomes 1,000 times, resulting in 1,000 data sets with randomized SCI-F values.

We randomized SCI-M values as described above for SCI-F, but with an additional control for patterns in sexual cycling (since sexual cycling attracts social attention from males; see Fig. S5) using data from Fig. S6 (randomization procedure visualized in Fig. S7). For all months after birth where a female had a live infant and had not resumed cycling, we substituted the matching SCI-M value from the randomly sampled female trajectory (i.e., representing the period when the randomly sampled female’s infant was alive and its mother had not yet resumed cycling). For all months after birth where a female had a live infant and had resumed sexual cycling, we substituted the SCI-M values from the randomly sampled trajectory during pre-pregnancy. For cases in which an infant died, we assigned the mother the randomized trajectory SCI-M value in pre-pregnancy, starting two months after the infant’s death and lasting for four months (reflecting rapid resumption of cycling in baboons after infant death and mean cycling length before the next conception: Zipple et al., 2017). After four months we assigned the pregnancy SCI-M value from the randomized trajectory.

We averaged the randomized monthly SCI values described above to get a mean SCI-F and SCI-M value for each infant’s mother during pregnancy, six months following birth, seven to 12 months following birth, and the shifting time window. We then ran parallel binomial GLMs on the randomized data sets to estimate the association between SCI measured over each time interval and infant survival, controlling for the same fixed effects as in analyses with observed data. The set of infant outcomes for each time interval matched the sample of outcomes for complementary analyses with real data in Tables S3 to S10.

After repeating this process across 1,000 randomized data sets, we found that in many cases the distribution of effect sizes generated from analyses with randomized data did not center on zero, indicating that infant-dependent and reproductive state-dependent changes in maternal social behavior indeed present a detectable confound over these time intervals (Fig. 3; Table S11). Furthermore, for both SCI-F and SCI-M, we found that all of the observed effect sizes fell well within the distribution of effect sizes generated from randomized data sets, indicating the observed effect sizes could be entirely explained by infant-dependent and reproductive state-dependent changes in maternal social behavior (Fig. 3; Table S11). In several cases the observed effects fell within the tails of the distributions (Fig. 3A, 3B, 3C, 3F, and 3G) making these effects less likely to be entirely explained by the confound than cases where effects fell within the center of the distribution. Among cases where the observed coefficients fell within one tail of a distribution, effects generated from SCI-F and SCI-M measured seven to 12 months after birth were the least likely to be entirely explained by the confound: less than one percent of coefficients from randomized data were larger than the observed coefficient (Fig. 3C and 3G; Table S11), in the direction of higher maternal sociality in this time interval being associated with *higher* infant mortality. Overall, these results suggest that infant-dependent and reproductive state-dependent patterns in maternal behavior are sufficient to produce non-zero associations between maternal sociality and infant survival when they are not taken into account, and that any evidence for maternal sociality having a consistent positive effect on infant survival is weak at best.

**Figure 3:**
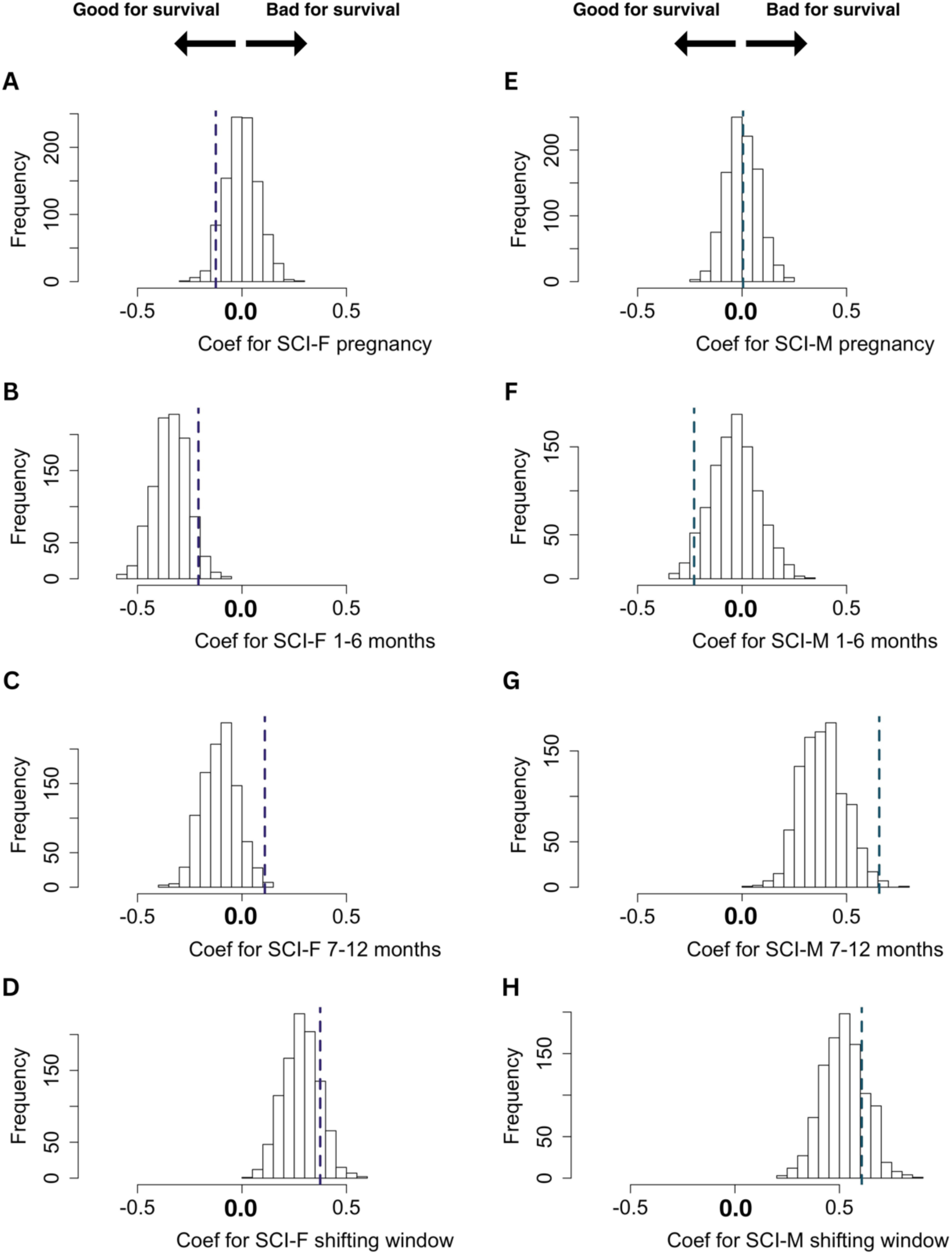
Results of the “Randomization of SCI Values” analysis. Histograms show the distribution of coefficients (i.e., the natural logarithm of the odds ratio) from binomial GLMs using randomized SCI data; vertical dashed lines show coefficients from GLMs with observed data (see Fig. 2). Left column shows SCI-F estimated over A) pregnancy, B) the six months following birth, C) the seven to 12 months following birth, and D) a shifting data window (six months prior to infant death or first birthday depending on survival outcome). Right column shows SCI-M estimated over E) pregnancy, F) the six months following birth, G) the seven to 12 months following birth, and H) a shifting data window. Negative coefficients mean high SCI is associated with low risk of mortality and thus high survival, while positive coefficients mean high SCI is associated with high risk of mortality and thus low survival. If an observed coefficient (dashed line) is to the *left* of the distribution of permuted coefficients, then high maternal sociality is associated with high infant survival after accounting for the confounding effect of infant and reproductive variation in maternal social behavior. If an observed coefficient is to the *right* of the distribution, then high maternal sociality is associated with high infant mortality after accounting for the confound. See Table S11 for median coefficient values and variances associated with distributions of coefficients from analyses on randomized data sets.

As a complementary test of this hypothesis, we conducted a second data randomization (hereafter ‘Randomization of Time Intervals’) that was designed to avoid (rather than quantify) any potential confound. Here, we randomly sampled, with replacement, the age at death for infants who died within one year, and assigned that age as the last day of the time interval used for SCI calculation for infants who survived. This approach avoids both the problem of sampling maternal behavior that occurs after an infant has died and the problem of measuring SCI at later ages for infants who survive. We then calculated maternal SCI-F and SCI-M over the six months prior to either the true death date (for dead infants) or the randomly sampled death date (for surviving infants). We analyzed data from 100 of these randomized data sets each with between 766 and 793 infant outcomes, depending on the number of infants whose mothers had complete behavioral data available for the sampled six month time interval, using GLMs that paralleled those applied above. These analyses provided no evidence that SCI-F or SCI-M predict infant survival (mean coef=−0.025; mean OR=0.975; 0% of p-values < 0.05 and mean coef=−0.029; mean OR=0.971; 0% of p-values < 0.05, respectively; distribution of coefficients shown in Fig. S8A and S8B).

### REPRODUCTIVE STATE AND INFANT-DEPENDENT TRENDS IN MATERNAL SOCIAL BEHAVIOR EXPLAIN THE EFFECT OF LIFETIME MATERNAL SOCIALITY ON INFANT SURVIVAL

The present analysis was motivated in part by previous results from Silk et al. (2003a), who showed that more socially integrated mothers experience higher infant survivorship. Silk et al. (2003a) used the individual mother (i.e., a female’s lifetime success at producing surviving infants) as the unit of analysis while the analyses reported above used the infant as the unit of analysis. Furthermore, Silk et al. (2003a) used a lifetime estimate of the “composite sociality index” (CSI), which combines data on female social relationships with both males and females (captured by grooming and proximity to others during focal points; see Supplementary Methods), as the measure of maternal sociality. This approach is in contrast to the measures we used above, which were summarized over short time intervals, sex-specific (SCI-F and SCI-M), and used only grooming data. These differences could mitigate the influence of the confound described above. To test this possibility, we recreated the analysis reported in Silk et al. (2003a) using an expanded data set that included an additional 22 years of behavioral and demographic data that have since accumulated. A total of 295 adult females were included in our recreation, compared to 108 in Silk et al. (2003a). See the Supplementary Methods for more details about our recreation.

We conducted three analyses on the expanded data set. First, following Silk et al. (2003a), we analyzed the relationship between a mother’s lifetime relative infant survival and her lifetime CSI (an estimate of her tendency to groom with and be in proximity to other adults). In this analysis, we recovered a similar pattern to the original paper: higher maternal sociality was associated with higher relative infant survival (B=0.095; p=0.027) (Fig. 4A), although the effect size was notably smaller than originally estimated (original paper reports B=0.321; p=0.015).

**Figure 4:**
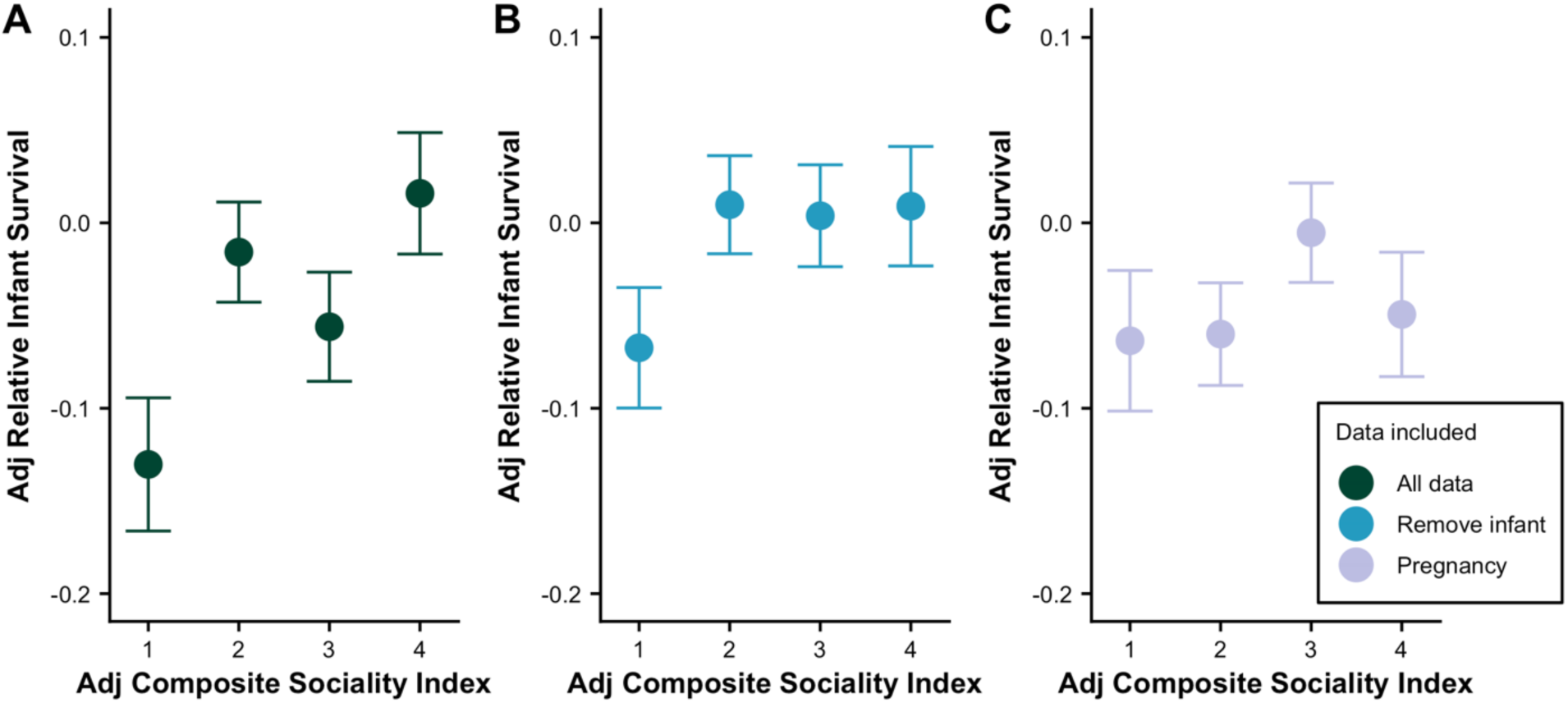
Recreation of Fig. 1 in Silk et al. (2003a) using an updated data set (1984 – 2022), showing the relationship between a mother’s lifetime Composite Sociality Index (CSI) score and relative infant survival. Panels from left to right show estimates: A) including all data from mothers’ adult lives (green), B) removing periods in the CSI calculation when mothers had a young infant less than one year old (blue), and C) only including data from periods of pregnancy (purple) in the CSI calculation.

Next, we repeated the analysis twice more, first removing time periods from the CSI calculation in which mothers had infants less than one year old, and second removing all time periods except pregnancy (when females typically do not have young infants and all share the same reproductive state). These two analyses limit the measures of maternal social behavior to periods when the mother’s behavior is least likely to be influenced by the presence of an attractive infant or by sexual cycling. In neither of these analyses did the relationship between maternal CSI and offspring mortality reach statistical significance (removing time periods where mothers had infants less than one year of age: B=0.031, p=0.268, n=256, Fig. 4B; restricting the analysis to periods of pregnancy: B=0.025, p=0.332, n=268, Fig. 4C). These results support the idea that the original result reported in Silk et al. (2003a) is primarily explained by infant-dependent and reproductive state-dependent trends in maternal social behavior.

### EFFECTS OF SOCIALITY ON ADULT SURVIVAL OUTWEIGH EFFECTS OF SOCIALITY ON OFFSPRING SURVIVAL

Our analysis revealed some limited evidence that mothers who are highly social during certain periods of their infant’s early life may experience modestly reduced offspring survival (Fig. 3C and 3G). At the same time, high SCI and DSI scores are associated with higher survival for adult female baboons themselves (Silk et al., 2010; Archie et al., 2014; Campos et al., 2020; Lange et al., 2023). If maternal sociality does indeed having conflicting influences on infant and maternal survival, how is the resulting tradeoff between maternal and infant survival resolved? To probe this question, we built a simple matrix projection model based on the life cycle of female baboons. The model’s parameters were chosen to roughly match the life history of our study population, and infant and adult female survival rates were assumed to be a function of SCI (males were not explicitly modeled; see Supplementary Methods for details).

Our primary goals were to determine what values of SCI maximize *λ*_1_—the leading eigenvalue from the transition matrix representing long-term per capita growth rate (the estimate of female fitness generated by the model; see Supplementary Methods)—and to understand how the relationship between SCI and *λ*_1_ changes depending on how SCI affects infant survival versus adult survival in our model. If *λ*_1_ is greatest when SCI is relatively high, this result would indicate that high levels of maternal sociality confer net fitness benefits for females. If *λ*_1_ is greatest when SCI is relatively low, this result would indicate that low levels of maternal sociality confer net fitness benefits (and conversely that high levels of maternal sociality impose net fitness costs).

We found that the effect of SCI on overall female fitness (*λ*_1_) depends almost entirely on how SCI affects adult female survival, regardless of the outcome for infants (Fig. 5). In Fig. 5, both of the top quadrants (where SCI is *positively* associated with adult survival) are almost entirely dark in color, indicating that high SCI *improves* fitness (*λ*_1_) when high SCI is good for adult survival (positive values on the y-axis), regardless of the effect of SCI on infant survival. Meanwhile both of the bottom two quadrants (where SCI is *negatively* associated with adult survival) are almost entirely light in color indicating that high SCI *reduces* fitness (*λ*_1_) when high SCI is bad for adult survival (negative values on the y-axis), regardless of the effect of SCI on infant survival. Notably, exceptions occur where the effects of SCI on adult survival are quite weak (i.e., the adult slope is close to zero): in these cases the infant slope becomes more influential in determining how SCI effects female fitness. The fact that SCI mainly influences fitness based upon its effects on adult survival likely follows from there being more adults in the population than infants at any given time.

**Figure 5:**
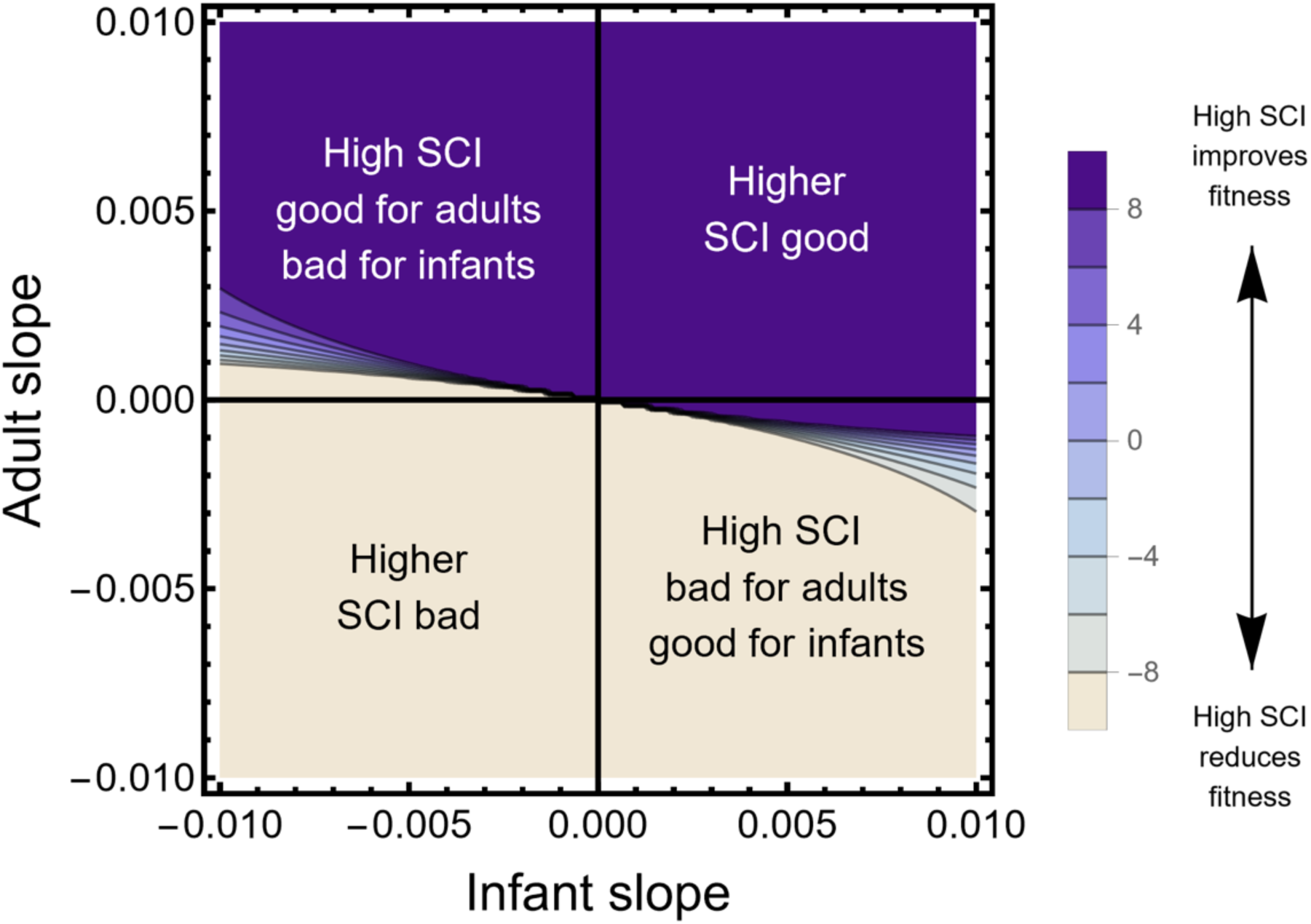
Colored landscape showing the value of SCI that maximizes *λ*_1_ (female fitness) in our matrix projection model. Dark colors correspond to cases where relatively high values of SCI improve female fitness, while light colors correspond to cases in which high values of SCI reduce female fitness (see color legend). Axes show the slope of infant survival (x-axis) and adult survival (x-axis) as a function of SCI when SCI = 0 (the population mean). More positive slopes correspond to a greater beneficial influence of high SCI on survival and more negative slopes correspond to a greater detrimental influence of high SCI on survival.

## DISCUSSION

### MATERNAL SOCIALITY AND INFANT SURVIVAL

We found no strong evidence that maternal sociality is causally associated with improved infant survival in our infant-wise analysis. We instead found that apparent relationships between maternal sociality and infant survival are more parsimoniously explained by infant-dependent and reproductive state-dependent trends in a mother’s social behavior. As a result, the direction of the relationship between maternal social behavior and infant survival changes depending on the time interval maternal sociality is captured over.

Our results further suggest that infant-dependent and reproductive state-dependent differences in maternal social behavior can generate a spurious positive relationship between maternal social relationships and infant survival not only in the first six months of an infant’s life, but even in the seven to 12 months following a birth. Specifically, Fig. 3C and Fig. 3G demonstrate that the distribution of effect sizes from analyses with randomized SCI-F and SCI-M data do not center on zero, indicating that removing data for the first 100 days following a birth (as in Silk et al., 2009) would not be enough to completely eliminate the effect of this confound in our population. Furthermore, controlling for this confound not only effects conclusions from infant-wise analyses, but also strongly attenuates the original effect reported in Silk et al. (2003a), which used the liftetime estimates of maternal sociality and infant survival (Fig. 4). Notably, when tested separately as predictors of relative infant survival in Silk et al. (2003a), proportion of time being groomed was the only one of the three social components of CSI that was a significant predictor of survival on its own (i.e., time spent grooming and spent in proximity to others are not significant predictors), suggesting that the amount of time mothers spend being groomed by others drives the relationship between CSI and infant survival. This further supports the idea that infants and certain reproductive states (e.g., sexual cycling) attracting social attention (expressed as directed grooming) contribute to the original result.

After accounting for variation in individual behavior driven by reproductive state and infants, our analyses with data from the Randomization of SCI Values analysis produced some limited evidence that more social mothers may experience higher infant mortality (Fig. 3C and 3G). A negative association between infant survival and maternal sociality captured over certain time periods could have multiple, non-mutually exclusive explanations. High levels of maternal social interactions could result in increased exposure to pathogens (May, 1983; Nunn, 2012), increased stress (Pearson et al., 2015), reduced time spent feeding (Altmann, 1980), or increased vulnerability to fatal kidnappings or infanticide (Kleindorfer & Wasser, 2004; Shopland & Altmann, 1987; Kalbitzer et al., 2017). Furthermore, a high frequency of social interactions between mothers and other females can result in rough handling of infants by other females, which in turn is associated with signs of distress in infants in Amboseli (Nguyen et al., 2009), and with lower survival in several other nonhuman primate populations (Silk 1980; Kleindorfer & Wasser 2004). On the other hand, maternal sociality may be negatively associated with infant survival if less healthy mothers or infants attract more social attention, for example, if kin and other close social partners intensify efforts to help support mothers when risk of offspring mortality appears heightened. In this case, maternal social bonds would not contribute directly to infant survival, but rather unhealthy mothers and infants would receive more social attention.

Notably, a potentially harmful effect of higher maternal sociality on infant survival was only supported over some time intervals used in our analyses. Analyses from our Randomization of Time Intervals approach did not support an effect of SCI on infant survival in either direction. Moreover, when SCI-F was measured over pregnancy and SCI-M was measured over the six months following a birth, the results of our Randomization of SCI Values were suggestive in the opposite direction: high SCI may be associated with slightly higher survival (although we could not rule out that these effects could be explained by a confound; Fig. 3A and 3F). The direction of these results with SCI-F are consistent with findings in Blersch et al. (2023) who found a positive relationship between maternal social connectedness and infant survival when measuring social relationships during pregnancy in an attempt to avoid the confound of infant attractiveness. Importantly, such a result could also be explained by reverse causality if, for example, mothers tend to socialize more when they are healthier (and thus more likely to birth a heathy infant). Regardless, given that we show maternal sociality has a neutral and occasionally even negative relationship with infant survival when quantified over other time periods, this suggests that mothers being more social does not provide universal fitness benefits for infants overall, at least in our population.

Furthermore, according to the results of our matrix projection model, any negative or positive effects of maternal sociality on infant survival are unlikely to shape maternal social behavior if maternal social relationships are linked to improved adult female survival. Specifically, even if maternal sociality directly affects both adult and infant outcomes, the survival benefits or costs that a mother experiences from being social outweigh the benefits or costs experienced by her infants. In other words, selection should not act to reduce sociality in baboon females if it improves their own survival, even if those social relationships pose potential risks to their infants. Consistent with adult survival having an outsized effect on fitness, findings suggest longevity is more important than fecundity for lifetime reproductive success in the Amboseli baboons (McLean et al., 2019).

## CONCLUSIONS

We have shown that the manner in which a female baboon’s social interactions are affected by her reproductive state and whether she has a live infant can lead to the erroneous inference that maternal sociality improves infant survival. Specifically, the manner in which a female’s social behavior changes depending upon her reproductive state and infant status generates a correlation between maternal sociality and infant survival that varies in both direction and magnitude depending on when sociality is measured relative to an infant’s birth or death (Fig. 3). Correcting for reproductive state and infant-related variation in maternal sociality can attenuate, eliminate, or even reverse the direction of maternal sociality-infant survival correlations. We believe that this confound is the best explanation for previous results reported from our study system, where high levels of maternal sociality were interpreted as a driver of enhanced infant survival (Silk et al., 2003a).

Taken together our results provide no strong support for a positive effect of maternal sociality on infant survival after accounting for this confound. In fact, the most compelling evidence for an association between maternal sociality and offspring survival was found in the seven to twelve months following birth where maternal sociality may be associated with poor infant survival in the Amboseli baboon population. This evidence that more social mothers may experience poor infant survival is suggestive rather than definitive: the strength of the relationship between high maternal sociality and poor infant survival varies across the time periods in which we measured maternal sociality (Fig. 3). Importantly, female state-dependent patterns of social interaction occur in other primate species (e.g., Gumert, 2007; Proctor et al., 2011; Tiddi et al., 2010), suggesting that our findings may generalize to other analyses. Consequently, our results mandate a closer scrutiny of the causal pathways linking maternal social behavior and infant survival in other nonhuman primate populations.

## METHODS

### ESTIMATING THE RELATIONSHIP BETWEEN INFANT SURVIVAL AND MATERNAL SOCIALITY, MEASURED OVER DIFFERENT TIME INTERVALS

#### Study system and subjects

Data for this analysis come from a long-term study of baboons (*Papio cynocephalus* and *P. anubis*) in the Amboseli basin of southern Kenya. This population has been under continuous observation since 1971 (Alberts & Altmann, 2012). All study subjects in our data set were live infants born into study groups between July of 1983 (when behavioral data began being collected systematically and consistently in our population) and December of 2020.

#### Maternal sociality

We quantified SCI (our measure of maternal sociality) in four six-month time intervals: (i) pregnancy, (ii) the fixed six-month window after the infant’s live birth (regardless of whether and when the infant died), (iii) the fixed six-month window beginning seven months after any infant’s live birth (regardless of whether and when the infant died), and (iv) a ‘shifting time window’ that represents the six months prior to each infant’s death or – if it survived – its first birthday. Maternal SCI with females (SCI-F) and with males (SCI-M) were calculated separately for each time interval using grooming data collected during “representative interaction sampling” (see Supplementary Methods) by averaging the residuals from two linear regressions: (i) log grooms given by the focal individual to other groupmates, regressed on log observer effort (i.e., the number of focal follows per adult female in the group divided by the number of days the female of interest was present in the group; see extended description in Supplementary Methods); and (ii) log grooms received by the focal individual, regressed on log observer effort (Archie et al., 2014). These values were z-scored within population and year and used as our final measure of SCI.

#### Testing the relationship between SCI and infant survival

We tested whether SCI-F or SCI-M predicted infant survival in separate models using binomial GLMs implemented with the glm function from R. Each model included one maternal social measure (SCI-F or SCI-M) measured over one of four time intervals. The outcome was scored according to whether the infant died (1) or survived (0). Given that the purpose of our analysis was to interrogate the previously reported linear effects of maternal sociality on infant survival and that no explicit hypotheses exist to support a non-linear relationship between sociality and infant survival, we did not explore the possibility of non-linear relationships in our analyses.

Several variables are already known or suspected to influence offspring survival in this population (Silk et al., 2003a; Zipple et al., 2021). To control for the possible influence of these variables on infant survival, we also measured them over the time intervals of interest and included them as additional fixed effects in our GLMs. These additional fixed effects were i) the relative proportion of adult females a given mother outranked in her group, ii) maternal age, (iii) maternal age squared, (iv) whether the infant of interest was the mother’s first birth (1=true; 0=false), and (v) group size. In models with SCI-M we also included the absolute proportion of days over the six-month time interval in which the mother was experiencing menstrual cycling as a fixed effect, to control for the ephemeral social relationships that arise between adult males and females when females are sexually cycling (Rowell, 1968; Seyfarth, 1978). In all models, continuous fixed effects were scaled by subtracting the mean and dividing by the standard deviation to ensure model convergence. We initially used mixed models including the random effects of maternal identity, hydrological year, and group identity in addition to fixed effects. However, these variables explained little to no variance leading to issues of singular fit and thus uninterpretable results. These terms were therefore dropped from all final analyses. Analyses of residuals with DHARMa (Hartig, 2017) indicated acceptable model fits.

## DATA AVAILABILITY STATEMENT

Data and code will be made available on Zenodo upon publication.

## ACKNOWLEDGEMENTS

We gratefully acknowledge the support of the National Science Foundation and the National Institutes of Health for the majority of the data represented here, most recently through R01AG53330, R01AG053308, R01AG071684, R01AG075914, and R61AG078470. M.J.A.C was supported by the Natural Sciences and Engineering Research Council of Canada. Support for ongoing data collection comes from the Max Planck Institute for Evolutionary Anthropology, and we thank Duke University, Princeton University, and the University of Notre Dame for financial and logistical support. We thank Steve Nowicki, Charles Nunn, Anne Pusey, Joseph Feldblum, and the Alberts lab group for their feedback on early versions of the manuscript and analysis. In Kenya, our research was approved by the Wildlife Research Training Institute, Kenya Wildlife Service, the National Commission for Science, Technology, and Innovation, and the National Environment Management Authority. We thank the University of Nairobi, the Kenya Institute of Primate Research, the National Museums of Kenya, the members of the Amboseli-Longido pastoralist communities, and the Enduimet Wildlife Management Area for their cooperation and assistance. We thank the Amboseli Baboon Project long-term field team (R.S. Mututua, S. Sayialel, J.K. Warutere, I.L., and Siodi, I.L.), and T. Wango and V. Oudu for their assistance in Nairobi. We thank N. Learn, J. Gordon, and W. Wilbur for database management. This research was approved by the IACUC at Duke University, the IACUC at the University of Notre Dame, and the Ethics Council of the Max Planck Society. The research adhered to all the laws and guidelines of Kenya. Visit http://amboselibaboons.nd.edu/acknowledgements/ for complete list of project acknowledgements.

