## Supplementary Materials for "Re-evaluating the relationship between female social bonds and infant survival in wild baboons"

### SUPPLEMENTARY TABLES

**Table S1. Comparison of SCI-F for females with surviving infants over four time intervals.**

Results of Tukey HSD test assessing the difference in mean SCI-F values for four time intervals using data from 257 female's with surviving infants (individual trajectories shown in Fig. 1A).

| Time intervals being compared | diff | lwr | upr | p adj |
|---|---|---|---|---|
| Pregnancy-Pre-pregnancy | 0.337 | 0.117 | 0.556 | <0.001** |
| Infant 1-6-Pre-pregnancy | 0.632 | 0.412 | 0.851 | <0.001** |
| Infant 7-12-Pre-pregnancy | 0.141 | -0.079 | 0.361 | 0.349 |
| Infant 1-6-Pregnancy | 0.295 | 0.075 | 0.515 | 0.003** |
| Infant 7-12-Pregnancy | -0.195 | -0.415 | 0.024 | 0.101 |
| Infant 7-12-Infant 1-6 | -0.491 | -0.710 | -0.271 | <0.001** |

+p≤0.1; * p<0.05; ** p<0.01.

**Table S2. Comparison of SCI-M for females with surviving infants over four time intervals.**

Results of Tukey HSD test assessing the difference in mean SCI-M values for four time intervals using data from 257 female's with surviving infants (individual trajectories shown in Fig. 1B).

| Time intervals being compared | diff | lwr | upr | p adj |
|---|---|---|---|---|
| Pregnancy-Pre-pregnancy | -0.879 | -1.074 | -0.685 | <0.001** |
| Infant 1-6-Pre-pregnancy | -0.973 | -1.168 | -0.779 | <0.001** |
| Infant 7-12-Pre-pregnancy | -1.202 | -1.397 | -1.007 | <0.001** |
| Infant 1-6-Pregnancy | -0.094 | -0.288 | 0.101 | 0.600 |
| Infant 7-12-Pregnancy | -0.322 | -0.517 | -0.128 | <0.001** |
| Infant 7-12-Infant 1-6 | -0.229 | -0.423 | -0.034 | 0.014* |

+p≤0.1; * p<0.05; ** p<0.01.

**Table S3. Maternal SCI-F as a predictor of infant survival, measured during pregnancy.**

Results from a binomial GLM that included other covariates (see text); n=923 infants from 273 mothers.

| | coefficient | OR | se | z | p |
|---|---|---|---|---|---|
| (Intercept) | -1.206 | 0.299 | 0.097 | -12.417 | <0.001** |
| SCI-F | -0.125 | 0.882 | 0.078 | -1.605 | 0.109 |
| Maternal proportional rank | -0.125 | 0.882 | 0.079 | -1.589 | 0.112 |
| Maternal age class | -0.813 | 0.444 | 0.556 | -1.463 | 0.144 |
| (Maternal age class)^2 | 0.984 | 2.675 | 0.514 | 1.916 | 0.055+ |
| First birth | 0.132 | 1.141 | 0.289 | 0.457 | 0.647 |
| Group size | -0.137 | 0.872 | 0.079 | -1.745 | 0.081+ |

Note: continuous variables were scaled by subtracting the mean and dividing by the standard deviation. +p≤0.1; * p<0.05; ** p<0.01.

**<u>Table S4.</u> Maternal SCI-F as a predictor of infant survival, measured in the six months after birth.**

Results from a binomial GLM that included other covariates (see text); n=857 infants from 263 mothers.

| | coefficient | OR | se | z | p |
|---|---|---|---|---|---|
| (Intercept) | -1.323 | 0.266 | 0.104 | -12.690 | <0.001** |
| SCI-F | -0.208 | 0.812 | 0.084 | -2.480 | 0.013* |
| Maternal proportional rank | -0.243 | 0.784 | 0.085 | -2.854 | 0.004** |
| Maternal age class | -1.385 | 0.250 | 0.622 | -2.226 | 0.026* |
| (Maternal age class)^2 | 1.493 | 4.451 | 0.577 | 2.589 | 0.010* |
| First birth | -0.109 | 0.897 | 0.310 | -0.351 | 0.726 |
| Group size | -0.218 | 0.804 | 0.086 | -2.537 | 0.011* |

Note: continuous variables were scaled by subtracting the mean and dividing by the standard deviation. +p≤0.1; * p<0.05; ** p<0.01.

**<u>Table S5.</u> Maternal SCI-F as a predictor of infant survival, measured in the seven to 12 months after birth.**

Results from a binomial GLM that included other covariates (see text); n=824 infants from 257 mothers.

| | coefficient | OR | se | z | p |
|---|---|---|---|---|---|
| (Intercept) | -1.499 | 0.223 | 0.112 | -13.406 | <0.001** |
| SCI-F | 0.109 | 1.116 | 0.093 | 1.174 | 0.240 |
| Maternal proportional rank | -0.248 | 0.780 | 0.092 | -2.681 | 0.007** |
| Maternal age class | -1.038 | 0.354 | 0.708 | -1.467 | 0.142 |
| (Maternal age class)^2 | 1.067 | 2.908 | 0.662 | 1.612 | 0.107 |
| First birth | -0.047 | 0.954 | 0.325 | -0.144 | 0.886 |
| Group size | -0.227 | 0.797 | 0.092 | -2.454 | 0.014* |

Note: continuous variables were scaled by subtracting the mean and dividing by the standard deviation. +p≤0.1; * p<0.05; ** p<0.01.

**<u>Table S6.</u> Maternal SCI-F as a predictor of infant survival, measured in a shifting time window.**

Results from a binomial GLM that included other covariates (see text); n=873 infants from 263 mothers.

| | coefficient | OR | se | z | p |
|---|---|---|---|---|---|
| (Intercept) | -1.171 | 0.310 | 0.101 | -11.644 | <0.001** |
| SCI-F | 0.374 | 1.454 | 0.092 | 4.075 | <0.001** |
| Maternal proportional rank | -0.314 | 0.731 | 0.086 | -3.657 | <0.001** |
| Maternal age class | -2.773 | 0.062 | 0.621 | -4.468 | <0.001** |
| (Maternal age class)^2 | 2.715 | 15.105 | 0.582 | 4.664 | <0.001** |
| First birth | -0.666 | 0.514 | 0.296 | -2.246 | 0.025* |
| Group size | -0.197 | 0.821 | 0.086 | -2.302 | 0.021* |

Note: continuous variables were scaled by subtracting the mean and dividing by the standard deviation. +p≤0.1; * p<0.05; ** p<0.01.

**<u>Table S7.</u> Maternal SCI-M as a predictor of infant survival, measured during pregnancy.**

Results from a binomial GLM that included other covariates (see text); ; n=923 infants from 273 mothers.

| | coefficient | OR | se | z | p |
|---|---|---|---|---|---|
| (Intercept) | -1.200 | 0.301 | 0.097 | -12.392 | <0.001** |
| SCI-M | 0.005 | 1.005 | 0.079 | 0.060 | 0.952 |
| Maternal proportional rank | -0.143 | 0.867 | 0.079 | -1.803 | 0.071+ |
| Maternal age class | -0.745 | 0.475 | 0.553 | -1.347 | 0.178 |
| (Maternal age class)^2 | 0.920 | 2.508 | 0.511 | 1.799 | 0.072+ |
| First birth | 0.117 | 1.124 | 0.288 | 0.405 | 0.686 |
| Group size | -0.127 | 0.881 | 0.078 | -1.615 | 0.106 |

Note: continuous variables were scaled by subtracting the mean and dividing by the standard deviation. +p≤0.1; * p<0.05; ** p<0.01.

**<u>Table S8.</u> Maternal SCI-M as a predictor of infant survival, measured in the six months after birth.**

Results from a binomial GLM that included other covariates (see text); n=857 infants from 263 mothers.

| | coefficient | OR | se | z | p |
|---|---|---|---|---|---|
| (Intercept) | -1.301 | 0.272 | 0.150 | -8.697 | <0.001** |
| SCI-M | -0.229 | 0.795 | 0.116 | -1.977 | 0.048* |
| Maternal proportional rank | -0.240 | 0.786 | 0.113 | -2.124 | 0.034* |
| Maternal age class | -1.065 | 0.345 | 0.793 | -1.342 | 0.180 |
| (Maternal age class)^2 | 1.325 | 3.764 | 0.724 | 1.831 | 0.067+ |
| First birth | -0.089 | 0.915 | 0.432 | -0.206 | 0.836 |
| Group size | -0.159 | 0.853 | 0.109 | -1.457 | 0.145 |
| Proportion of days mother was cycling | 2.447 | 11.555 | 0.273 | 8.950 | <0.001** |

Note: continuous variables were scaled by subtracting the mean and dividing by the standard deviation. +p≤0.1; * p<0.05; ** p<0.01.

**<u>Table S9.</u> Maternal SCI-M as a predictor of infant survival, measured in the seven to 12 months after birth.**

Results from a binomial GLM that included other covariates (see text); n=824 infants from 257 mothers.

| | coefficient | OR | se | z | p |
|---|---|---|---|---|---|
| (Intercept) | -1.585 | 0.205 | 0.118 | -13.422 | <0.001** |
| SCI-M | 0.656 | 1.928 | 0.119 | 5.521 | <0.001** |
| Maternal proportional rank | -0.315 | 0.730 | 0.095 | -3.316 | 0.001** |
| Maternal age class | -1.241 | 0.289 | 0.719 | -1.726 | 0.084+ |
| (Maternal age class)^2 | 1.238 | 3.450 | 0.672 | 1.842 | 0.065+ |
| First birth | -0.037 | 0.963 | 0.332 | -0.113 | 0.910 |
| Group size | -0.233 | 0.792 | 0.095 | -2.442 | 0.015* |
| Proportion of days mother was cycling | -0.304 | 0.738 | 0.112 | -2.708 | 0.007** |

Note: continuous variables were scaled by subtracting the mean and dividing by the standard deviation. +p≤0.1; * p<0.05; ** p<0.01.

**Table S10.** **Maternal SCI-M as a predictor of infant survival, measured in a shifting time window.**

Results from a binomial GLM that included other covariates (see text); n=873 infants from 263 mothers.

| | coefficient | OR | se | z | p |
|---|---|---|---|---|---|
| (Intercept) | -1.831 | 0.160 | 0.168 | -10.928 | <0.001** |
| SCI-M | 0.607 | 1.834 | 0.111 | 5.446 | <0.001** |
| Maternal proportional rank | -0.146 | 0.864 | 0.096 | -1.524 | 0.128 |
| Maternal age class | -3.156 | 0.043 | 0.700 | -4.512 | <0.001** |
| (Maternal age class)^2 | 3.016 | 20.419 | 0.660 | 4.573 | <0.001** |
| First birth | -0.913 | 0.401 | 0.328 | -2.782 | 0.005** |
| Group size | -0.383 | 0.682 | 0.102 | -3.770 | <0.001** |
| Proportion of days mother was cycling | -2.247 | 0.106 | 0.251 | -8.945 | <0.001** |

Note: continuous variables were scaled by subtracting the mean and dividing by the standard deviation. +p≤0.1; * p<0.05; ** p<0.01.

**Table S11.** **Model results for data sets generated via the Randomization of SCI Values.**

Distribution of coefficients from binomial GLMs testing the relationship between randomized SCI-F and randomized SCI-M, and infant survival. Each row describes the distribution of coefficients from 1000 randomized data sets.

| measure of maternal sociality | time window | median coefficient from randomized data | variance of distribution of coefficients | % above observed coefficient | % below observed coefficient |
|---|---|---|---|---|---|
| SCI-F | Pregnancy | -0.001 | 0.006 | 95.3 | 4.7 |
| | 1-6 m after birth | -0.337 | 0.007 | 5.5 | 94.5 |
| | 7-12 m after birth | -0.103 | 0.007 | 0.6 | 99.4 |
| | Shifting time window | 0.286 | 0.007 | 14.6 | 85.4 |
| SCI-M | Pregnancy | -0.002 | 0.006 | 47.0 | 53.0 |
| | 1-6 m after birth | -0.039 | 0.013 | 95.7 | 4.3 |
| | 7-12 m after birth | 0.381 | 0.011 | 0.7 | 99.3 |
| | Shifting time window | 0.520 | 0.010 | 20.3 | 79.7 |

### SUPPLEMENTARY FIGURES

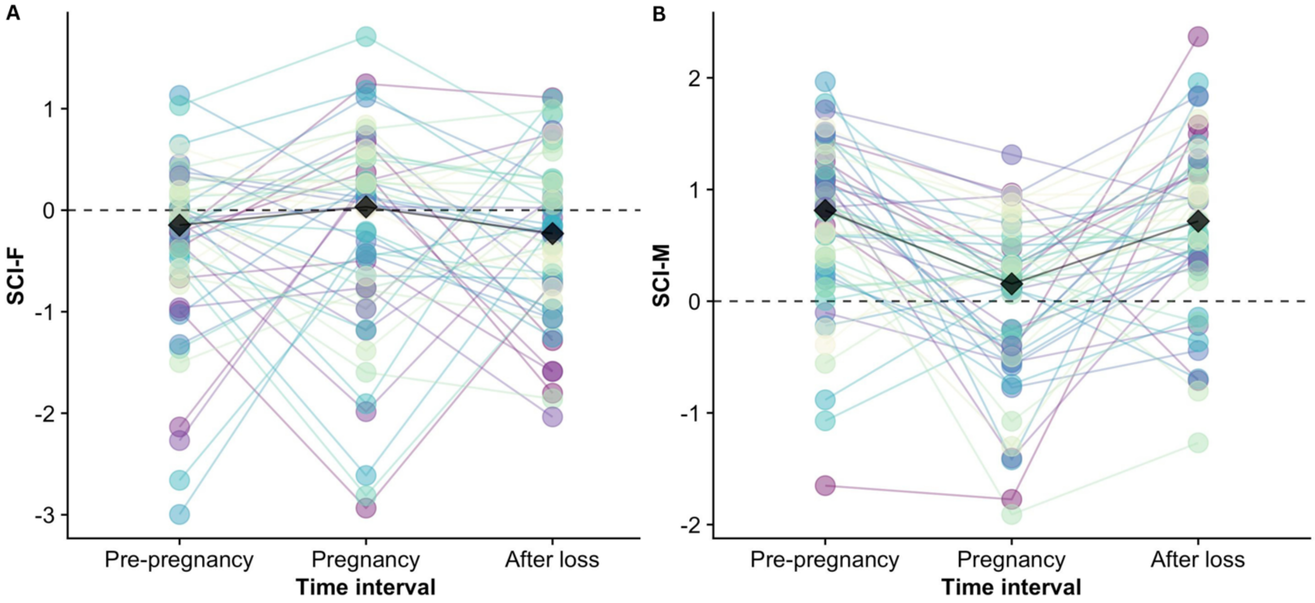

**Figure S1. Comparison of A) social connectedness to females (SCI-F) and B) social connectedness to males (SCI-M) for mothers with infants who died in the first year of life.**

Values are shown across three six-month time interval (six months prior to pregnancy, pregnancy, and after the loss of the infant i.e., the six months following the infant's death). Black diamonds show the median for each time interval. Each series of connected dots represents a single infant ID for 51 unique infant births (n=45 unique mothers) where the infant died within the first six months after birth, behavioral data for the mother was available for all three time intervals, and the mother did not have a young infant (less than six months old) from a previous pregnancy during the pre-pregnancy period.

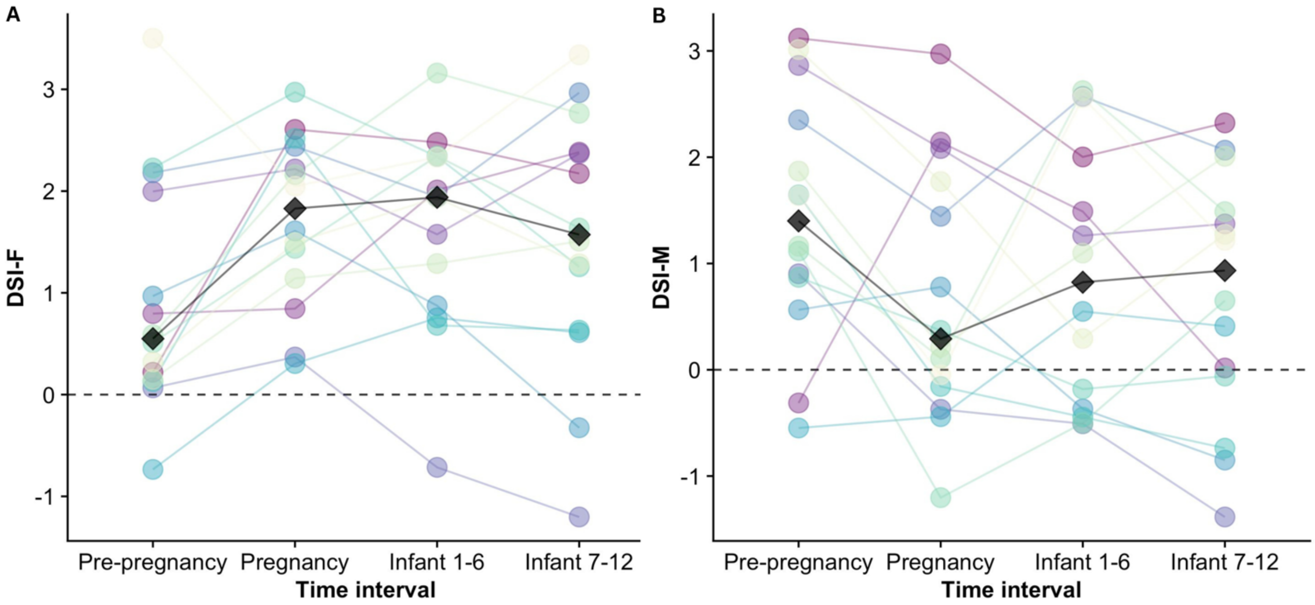

**<u>Figure S2.</u> Comparison of scores on A) the dyadic sociality index with females and B) the dyadic sociality index with males for mothers with infants who survived to one year.**

Values are shows across four six-month time intervals (six months prior to pregnancy, pregnancy, one to six months following birth, and seven to 12 months following birth). Black diamonds show the median for each time interval. Color encodes infant ID. Sample of 14 infant births was used to reduce computational demands. All mothers had behavioral data available for all four time intervals and did not have a young infant (less than six months old) from a previous pregnancy during the pre-pregnancy period.

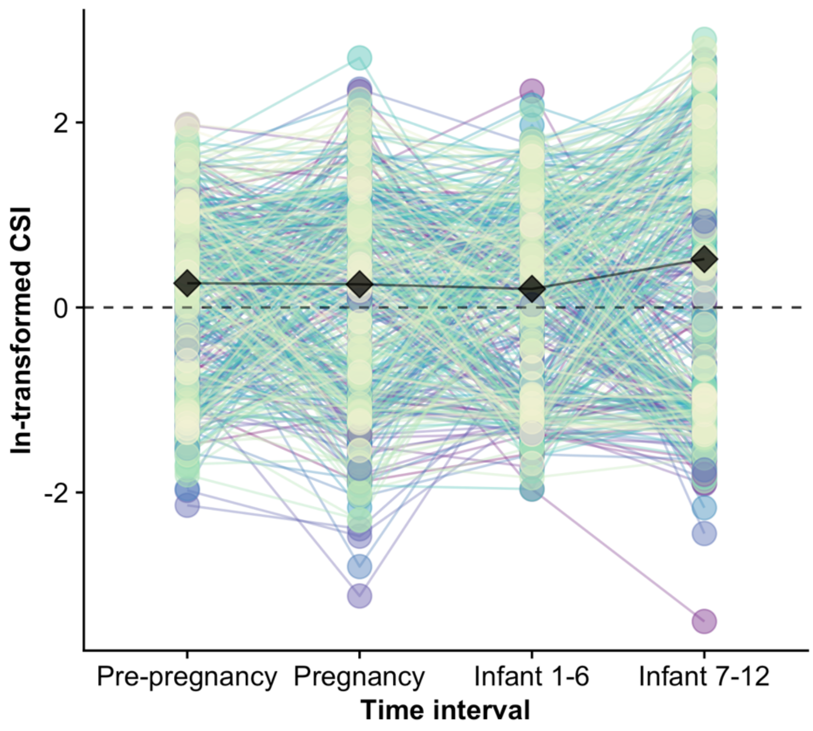

**<u>Figure S3.</u> Comparison of composite sociality index (CSI, which includes social interactions with both sexes) scores for mothers with infants who survived to one year.**

Values are ln-transformed scores shown across four six-month time intervals (six months prior to pregnancy, pregnancy, one to six months following birth, and seven to 12 months following birth). Black diamonds show the median for each time interval. Each series of connected dots represents a single infant ID for 418 unique infant births (n=198 unique mothers) where behavioral data for the mother was available for all four time intervals and the mother did not have a young infant (less than six months old) from a previous pregnancy during the pre-pregnancy period. Because CSI is calculated using data from focal samples (see Supplementary Methods), only infants whose mothers had more than 50 focal points taken during each time interval were included in this figure.

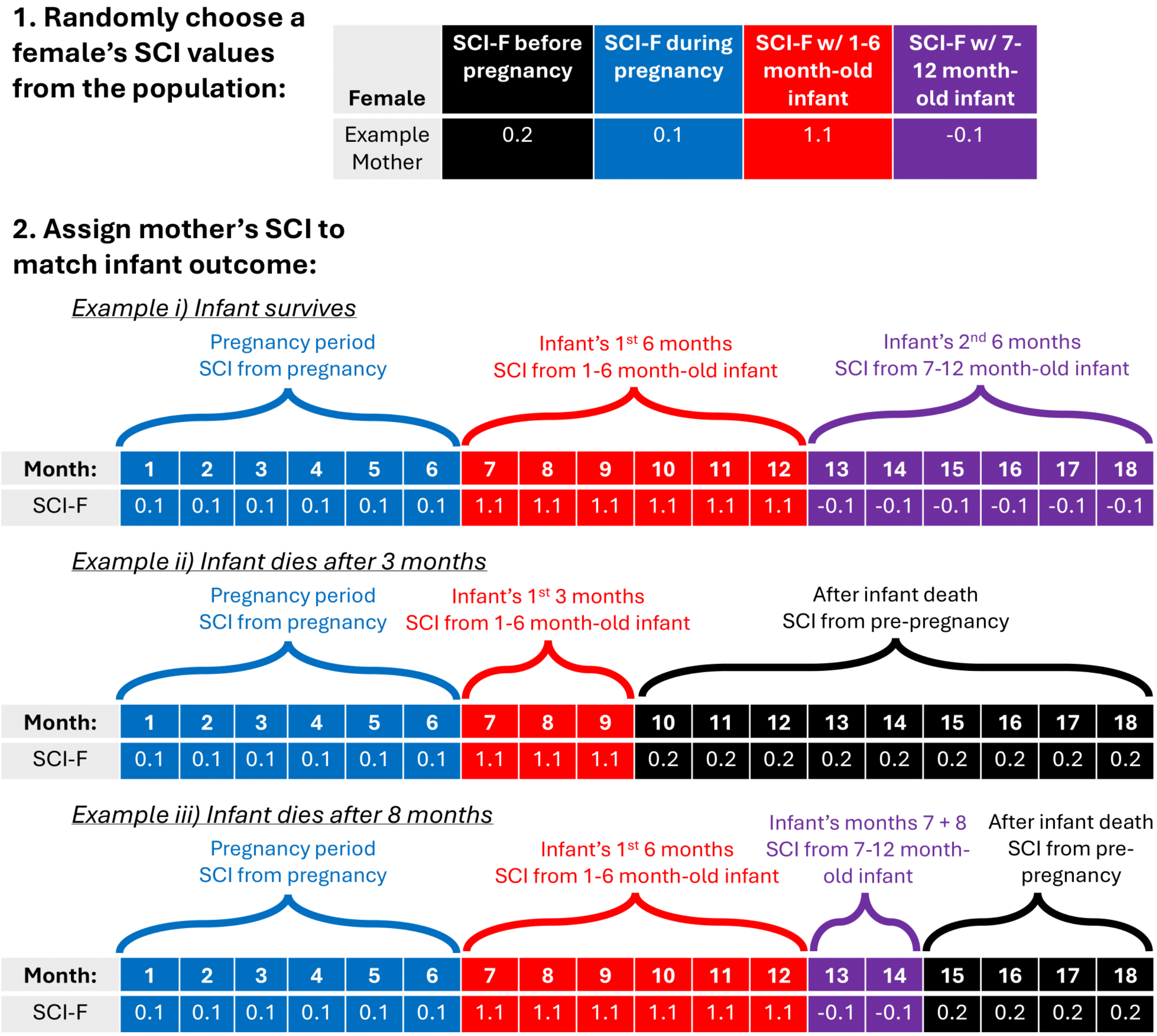

**Figure S4. Visualization of the Randomization of SCI Values approach for SCI-F.**

Randomization process for SCI-F is described in text. For each mother-infant pair in the real data, 1: Choose a trajectory of SCI-F values from a random mother with a surviving infant shown in Fig. 1. The second row in the table shows the values of SCI-F from the randomly selected mother in the four time intervals. 2: Assign SCI-F values for each time interval, depending on the infant outcome in the real data.

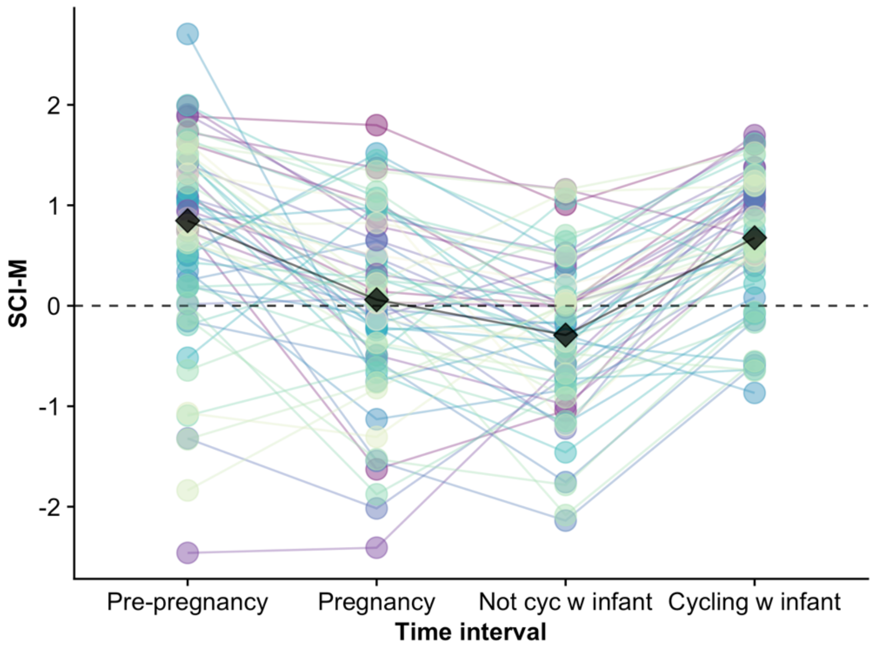

**<u>Figure S5.</u> Comparison of social connectedness to males (SCI-M) for mothers with infants who survived to one year including after cycling resumption.**

Values are shown across four time intervals (six months prior to pregnancy, pregnancy, the time a mother spent with the live infant before resuming cycling, and the time a mother spent with the live infant after resuming cycling). Black diamonds show the median for each time interval. Each series of connected dots represents a single infant ID for 59 unique infant births (n=46 unique mothers) where behavioral data for the mother was available for all four time intervals, where the mother resumed cycling at least 90 days before the infant's first birthday (to ensure enough behavioral data were available during that period to calculate SCI-M), and where the mother did not have a young infant (less than six months old) from a previous pregnancy during the pre-pregnancy period.

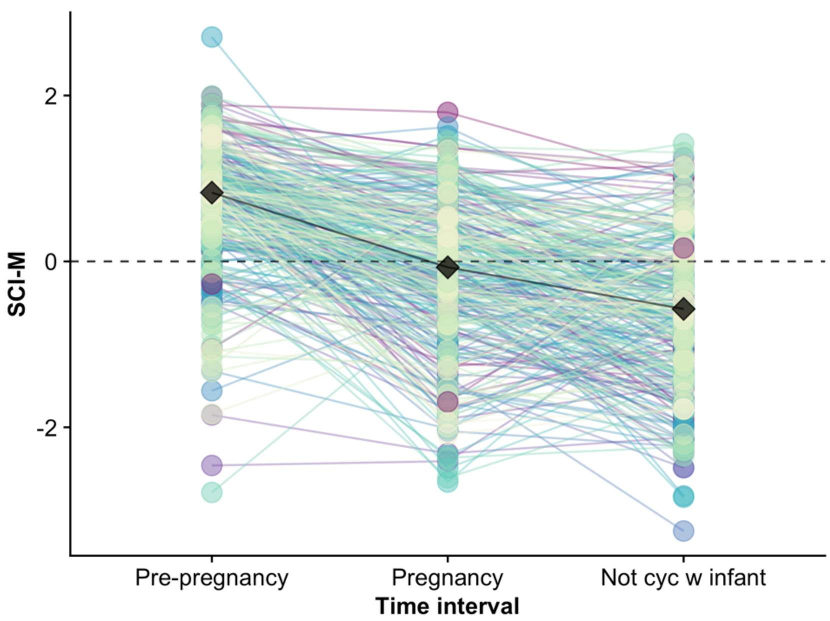

**Figure S6. Comparison of social connectedness to males (SCI-M) for mothers with infants who survived to one year.**

Values are shown across three time intervals (six months prior to pregnancy, pregnancy, and the time a mother spent with the live infant before resuming cycling). Black diamonds show the median for each time interval. Each series of connected dots represents a single infant ID for 313 unique infant births (n=165 unique mothers) where behavioral data for the mother was available for all three time intervals and the mother did not have a young infant (less than six months old) from a previous pregnancy during the pre-pregnancy period.

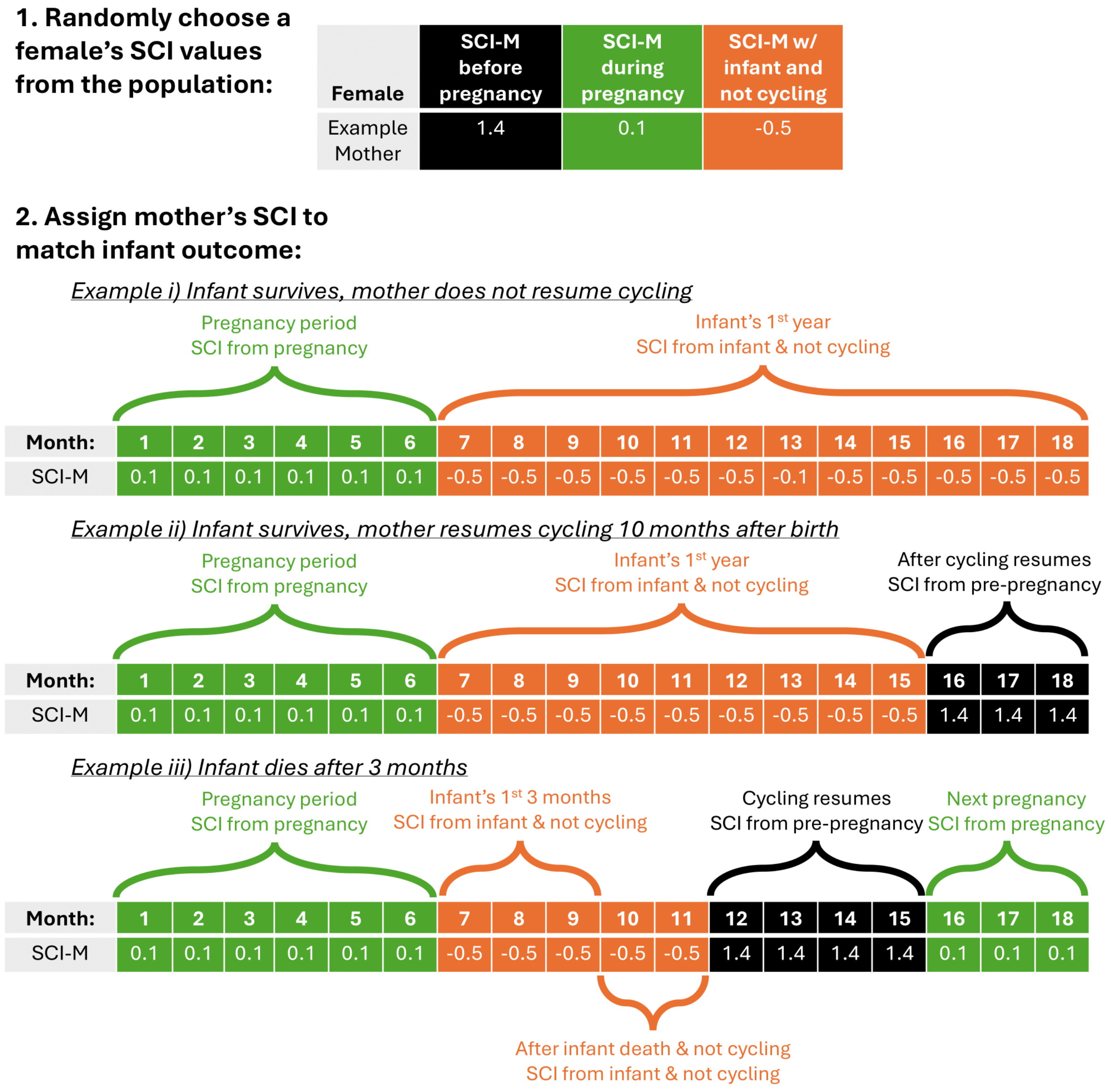

**Figure S7. Visualization of the Randomization of SCI Values approach for SCI-M.** Randomization process for SCI-M is described in text. For each mother-infant pair in the real data, 1: Choose a trajectory of SCI-M values from a random mother with a surviving infant shown in Fig. S4. The second row in the table shows the values of SCI-M from the randomly selected mother in the four time intervals. 2: Assign SCI-M values for each time interval, depending on the infant outcome in the real data.

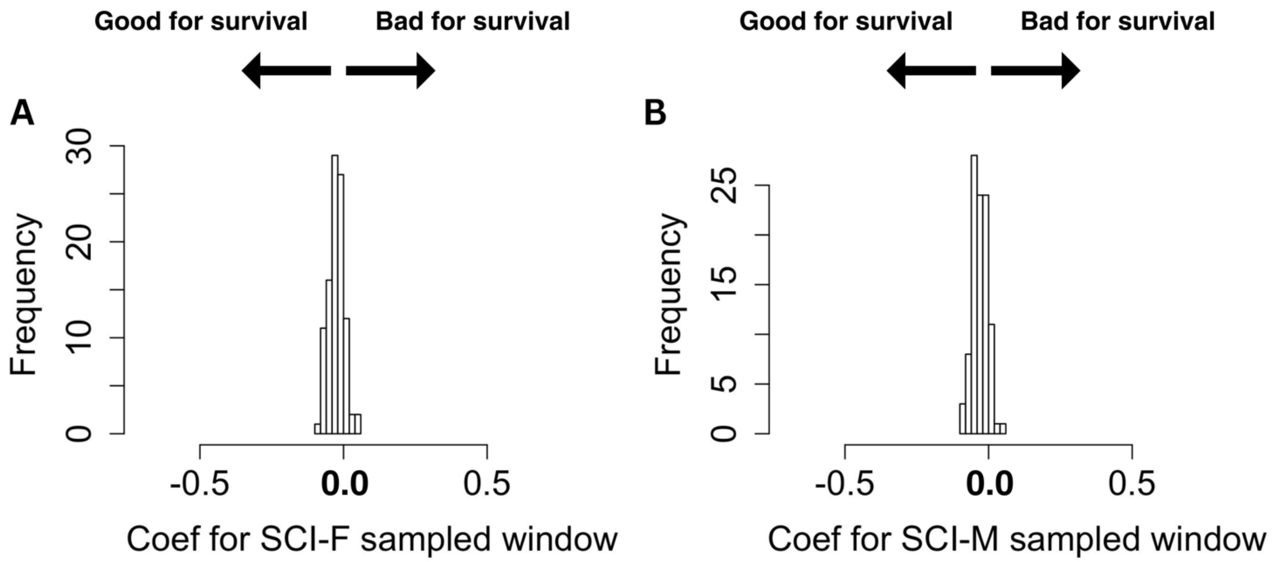

**<u>Figure S8.</u> Histograms of coefficients showing the effect of maternal sociality on infant survival when maternal sociality is measured using the Randomization of Time Intervals approach.** Coefficients are from 100 binomial GLMs each with a sample size of between 766 and 793 infant outcomes where maternal A) SCI-F and B) SCI-M are measured over the six months prior to the death date of infants who died in the first year and six months prior to a randomly sampled dead infant's age at death for infants who survived to one year. For both SCI-F and SCI-M, the coefficients center on zero suggesting no effect.

### **SUPPLEMENTARY METHODS**

#### CALCULATING OBSERVER EFFORT FOR SCI

Observers recorded dyadic grooming as part of regular “representative interaction sampling”: during 10-minute long focal samples, observers record all grooming interactions between any group members in their line of sight. During focal animal sampling, the observers move from subject to subject in a randomized order, ensuring representative sampling of the social group. However, estimating interaction rates from the representative interaction sampling data is complicated by the fact that the number of observers remained constant while group sizes varied, resulting in higher numbers of grooming interactions per dyad recoreded in smaller groups than in larger ones. We calculated observer effort by taking the total number of focal follows completed in a group during the time-period of interest divided by the mean number of females present in the group across all days in the study period. This number was then further divided by the number of days the female of interest was present in the group. The resulting value was our measure of observer effort used to calculate SCI.

#### CALCULATING DSI AND CSI

To assess whether the trends in maternal social behaviors captured by SCI were similar using other indexes of maternal sociality, we also looked at trends in two other measures of maternal sociality: i) the dyadic sociality index (DSI) — a sex-specific measure of the strength of an individual’s top three bonds, and ii) the composite sociality index (CSI) — the measure of sociality used in the Silk et al. (2003) lifetime analysis, which considers the social relationships of the focal individual with adults of both sexes in one combined metric.

To calculate DSI for mothers during each of the four time intervals (six months prior to pregnancy, pregnancy, one to six months following birth, and seven to 12 months following birth) we followed the methods used in Campos et al. (2021): i) we calculated relative daily grooming rates for every *dyad* in the population between the dates matching to each mother’s pre-pregnancy period, pregnancy period, one to six months following birth, and seven to 12 months following birth as the female’s number of grooming interactions (in either direction) divided by the number of co-residence days for that dyad during the relevant time window; ii) we calculated observer effort during each dyad’s co-residence days in a given group and year as the number of focal samples on adult females collected during those days, divided by the mean number of adult females in the group during those days, divided by the number of co-residence days for that dyad; iii) we then calculated an index of dyadic bond strength by taking the residuals of a regression between log daily grooming rate against log observer effort; iv) we calculated the two dyadic sociality indices for each mother, bond strength with females (DSI-F) and bond strength with males (DSI-M), as the mean of the dyadic bond strength values for her three strongest adult female and adult male grooming partners, respectively.

CSI was calculated over each six-month time interval per Silk et al. (2003) using data from focal samples.

$$CSI = \frac{\frac{prop\ grooms\ given}{median\ prop\ given} + \frac{prop\ grooms\ received}{median\ prop\ received} + \frac{prop\ proximity}{median\ prop\ proximity}}{3}. \quad (S1)$$

RECREATION OF SILK ET AL. (2003)

To recreate the analysis documented in Silk et al. (2003) using the expanded data set now available for the Amboseli baboons, we estimated CSI over the entire adult period for 295 adult females who had infants before 2022 following the methods outlined in Silk et al. (2003). Over the same adult period, we estimated each female's relative infant survival: the difference between the proportion of a mother's infants that survived to one year of age and the median proportion of all infants from the population that survived to one year. This measure included both live births and fetal losses per the original study.

We corrected both maternal CSI and relative infant survival for a shift in habitat use by our study groups as done in Silk et al. (2003). This home range shift was associated with striking changes in baboon behavior, demography, and DNA methylation patterns (Silk et al., 2003; Alberts et al., 2005; Anderson et al., 2024). Following Silk et al. (2003) we created adjusted values by dividing our data set into "pre-move" and "post-move" time-periods. The pre-move time-period was further separated by group, since the two study groups from that time differed in when they transitioned habitats (Bronikowski & Altmann, 1996). We calculated CSI scores for females present during the "pre-move" and "post-move" time-periods separately using the CSI equation above (S1) and using time-period specific (and when relevant group specific) pre- and post- move medians for proximity, grooms given, and grooms received. Adjusted relative infant survival was estimated as the difference between a female's own proportion of surviving infants and the proportion of all infants that survived during the time-period of interest. If a female was observed in both the pre- and post- move time periods, we assigned them a CSI and relative infant survival score that was the average of their pre- and post- move scores.

We tested the relationship between habitat-corrected lifetime CSI and habitat-corrected relative infant survival in a linear regression, including mean female dominance rank as an additional fixed effect. Note that this model structure and analysis procedure differs from the GLM and covariate set included in the primary analyses in this paper, in order to be as consistent as possible with the analysis conducted by Silk et al. (2003).

Finally, to assess how infant-dependent patterns in social behavior contributed to these results, we repeated this analysis using two separate approaches. First, we repeated the analysis but removed focal data from CSI calculations that occurred within one year of an infant's birth. Second, we repeated the analysis but removed all focal data except those collected when a female was pregnant.

<u>MATRIX PROJECTION MODEL</u>

We treated survival of females as a function of SCI, denoted by $x$, with the form

$$s(\mu, c, x) = \mu^{\exp(-c\,x)}. \quad \text{(S2)}$$

Here, $\mu$ is mean survival rate when $x = 0$ and $c$ is a coefficient describing how survival changes with SCI $x$. In particular, when $c > 0$, survival decreases with increasing SCI; when $c < 0$, survival increases with increasing SCI. A larger magnitude of $c$ corresponds to a stronger influence of SCI on survival. It is worth noting that the absolute influence of SCI $x$ on survival $s$ assumed by equation (S2) depends on both $\mu$ and $c$, such that:

$$\left.\frac{ds}{dx}\right|_{x=0} = c\,\mu\log(\mu). \quad \text{(S3)}$$

Thus, rather than focusing on the parameters $\mu$ and $c$ directly, we used equation (S3), the slope of the survival probability as a function of SCI $x$ when $x = 0$, as a measurement of the effect of SCI on survival (see, e.g., Fig. 5).

Model dynamics were in discrete time and are visualized in Fig. S9A. A proportion $s_I = s(\mu_I, c_I, x)$ of infants survive from time step $t$ to time step $t + 1$ to become first year juveniles. Likewise, a proportion of $s_J = s(\mu_J, 0, x)$ juveniles survive. Note that we assumed that juvenile survival is independent of maternal SCI to isolate the influence of SCI on infant and adult survival. Of surviving juveniles, a proportion $m$ mature to become adults. Adults survive and remain in the adult stage with probability $s_A = s(\mu_A, c_A, x)$. Each adult produces $f$ offspring each time step. We set mean survival, maturation, and fecundity to loosely match the pattern from the study population (i.e., that of a long-lived organism with higher mortality early and late in life). Figure S9B lists all parameter values, with $c_J$ and $c_A$ varying throughout our analyses.

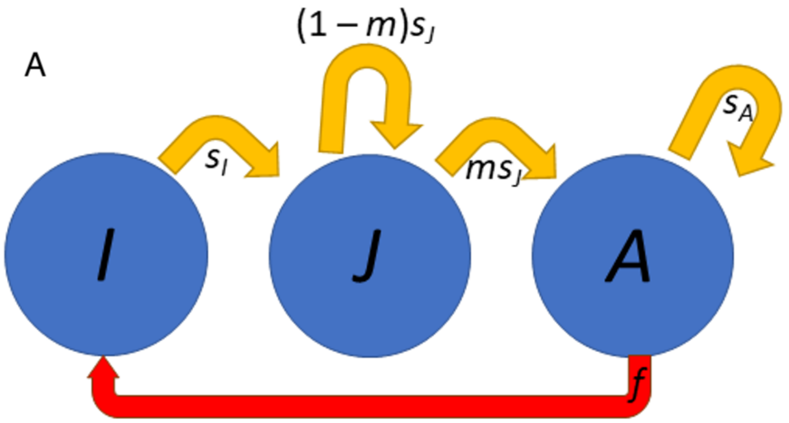

B

| Symbol | Meaning | Value |
|---|---|---|
| $\mu_I$ | Mean infant survival | 0.75 |
| $\mu_J$ | Mean juvenile survival | 0.9 |
| $\mu_A$ | Mean adult survival | 0.95 |
| $m$ | Proportion maturing | 0.3 |
| $f$ | Fecundity | 0.25 |

**<u>Figure S9.</u> Overview of matrix projection model.**
A) Flow diagram demonstrating transitions between life stages. We used a three-state model: infants, juveniles, and adults (labeled I, J, and A, respectively). B) Parameter values used throughout.

The dynamics of matrix projection models can be concisely written as

$$\vec{n}(t+1) = M\vec{n}(t). \qquad \text{(S4)}$$

Here, $\vec{n} = (I \quad J \quad A)^T$ is the vector of densities in the various stages. $M$ is known as the "transition matrix" and describes transitions between the stages. It is well known that the eigensystem of the transition matrix $M$ is key in determining the long-run dynamics of linear dynamical systems (Caswell, 2000). In particular, the system (equation S4) will converge to the stage distribution given by the right eigenvector, $v_1$, associated with the largest eigenvalue (in terms of magnitude) of $M$, $\lambda_1$. For this model,

$$M = \begin{pmatrix} 0 & 0 & f \\ s_I & (1-m)s_J & 0 \\ 0 & ms_j & s_A \end{pmatrix}. \tag{S5}$$

The leading eigenvalue of $M$, $\lambda_1$, gives the long-run growth rate of the system (i.e., population mean fitness).

The assumptions of our model mean that the leading eigenvalue $\lambda_1$ has both ecological and evolutionary implications. Ecologically, $\lambda_1$ is simply the long-run per capita growth rate, and thus increasing $\lambda_1$ corresponds to a population-level benefit. However, SCI $x$ is a parameter of the matrix projection model (equation S4). In other words, the simple matrix projection model is best interpreted as a population in which each individual has the same SCI. Consider, however, what would happen if a "mutant" strategy with a different SCI was introduced in the population. In the absence of frequency dependence, we can simply compare the leading eigenvalue of the initial population with that of a population consisting only of mutants $\lambda_1'$. If $\lambda_1' > \lambda_1$, then the mutant strategy will grow geometrically faster (or shrink geometrically slower) than the resident population, and thus, given sufficiently long time, become the dominant strategy and take over the population (i.e., the frequency of the mutant strategy will approach one). Conversely, if $\lambda_1' < \lambda_1$, then the mutant strategy will remain exceedingly rare indefinitely. This simple argument conveys why $\lambda_1$ can also be viewed as a measure of fitness (specifically, invasion fitness; Caswell, 2000). Strategies leading to a higher $\lambda_1$ can be expected to become more common under the simplifying assumptions of our model, and thus values of SCI that maximize $\lambda_1$ are uninvasible "evolutionarily stable strategies" (i.e., evolutionary endpoints).

Our initial analysis was to determine what values of SCI $x$ maximize $\lambda_1$. We varied the strength of the trade-off between infant and adult survival by changing the coefficients $c_I$ and $c_A$. As seen in Fig. 5, rather than reporting $c_J$ and $c_A$ directly, we focus on the slope of survival as a function of SCI (equation S3) to aid in interpretation. Biologically, we seek the values of SCI that lead to the fastest per capita growth rate and that would also be favored by selection.

A few important caveats apply to this model. First, while SCI is in practice calculated for each focal individual relative to the rest of the population (i.e., it is a *relative* measure of sociality, so it is impossible for all individuals to increase their SCI values), our projection model treats the parameter $x$ as monomorphic (i.e., it is an *absolute* measure, making it possible for the whole population to become "more" or "less" socially integrated). In other words, there is some incompatibility with how SCI is measured in the Amboseli baboons in practice and how our model can be interpreted: if the fitness consequences of sociality are only or primarily absolute (rather than a consequence of sociality relative to conspecifics), our projection analysis will be most applicable to the study population. If, however, the fitness consequences of sociality are only or primarily relative, its applicability is limited. In nature, there are likely both relative and absolute fitness consequences of sociality. Second, this model is phenomenological, that is it describes the empirical relationship of one variable to another, and does not include any mechanism by which sociality influences survival. Thus, we do not intend to use the model to make conclusions about SCI per se, but rather intened to draw conclusions about an arbitrary trait that has contrasting influences on adult versus infant survival.
